# Using machine learning to guide targeted and locally-tailored empiric antibiotic prescribing in a children’s hospital in Cambodia

**DOI:** 10.1101/367037

**Authors:** Mathupanee Oonsivilai, Mo Yin, Nantasit Luangasanatip, Yoel Lubell, Thyl Miliya, Pisey Tan, Lorn Loeuk, Paul Turner, Ben S Cooper

## Abstract

**Background:** Early and appropriate empiric antibiotic treatment of patients suspected of having sepsis is associated with reduced mortality. The increasing prevalence of antimicrobial resistance risks eroding the benefits of such empiric therapy. This problem is particularly severe for children in developing country settings. We hypothesized that by applying machine learning approaches to readily collected patient data, it would be possible to obtain actionable and patient-specific predictions for antibiotic-susceptibility. If sufficient discriminatory power can be achieved, such predictions could lead to substantial improvements in the chances of choosing an appropriate antibiotic for empiric therapy, while minimizing the risk of increased selection for resistance due to use of antibiotics usually held in reserve.

**Methods and Findings:** We analyzed blood culture data collected from a 100-bed children’s hospital in North-West Cambodia between February 2013 and January 2016. Clinical, demographic and living condition information for each child was captured with 35 independent variables. Using these variables, we used a suite of machine learning algorithms to predict Gram stains and whether bacterial pathogens could be treated with standard empiric antibiotic therapies: i) ampicillin and gentamicin; ii) ceftriaxone; iii) at least one of the above.

243 cases of bloodstream infection were available for analysis. We used 195 (80%) to train the algorithms, and 48 (20%) for evaluation. We found that the random forest method had the best predictive performance overall as assessed by the area under the receiver operating characteristic curve (AUC), though support vector machine with radial kernel had similar performance for predicting Gram stain and ceftriaxone susceptibility. Predictive performance of logistic regression, simple and boosted decision trees and k-nearest neighbors were poor in comparison. The random forest method gave an AUC of 0.91 (95%CI 0.81-1.00) for predicting susceptibility to ceftriaxone, 0.75 (0.60-0.90) for susceptibility to ampicillin and gentamicin, 0.76 (0.59-0.93) for susceptibility to neither, and 0.69 (0.53-0.85) for Gram stain result. The most important variables for predicting susceptibility were time from admission to blood culture, patient age, hospital versus community-acquired infection, and age-adjusted weight score.

**Conclusions:** Applying machine learning algorithms to patient data that are readily available even in resource-limited hospital settings can provide highly informative predictions on susceptibilities of pathogens to guide appropriate empiric antibiotic therapy. Used as a decision support tool, such approaches have the potential to lead to better targeting of empiric therapy, improve patient outcomes and reduce the burden of antimicrobial resistance.

**Author summary:**

Why was this study done?
- Early and appropriate antibiotic treatment of patients with life-threatening bacterial infections is thought to reduce the risk of mortality.
- In hospitals that have a microbiology laboratory, it takes 3-4 days to get results which indicate which antibiotics are likely to be effective; before this information is available antibiotics have to be prescribed *empirically* i.e. without knowledge of the causative organism.
- Increasing resistance to antibiotics amongst bacteria makes finding an appropriate antibiotic to use empirically difficult; this problem is particularly severe for children in developing country settings.
- If we could predict which antibiotics were likely to be effective at the time of starting antibiotic therapy, we might be able to improve patient outcomes and reduce resistance.

What Did the Researchers Do and Find?
- We evaluated the ability of a number of different algorithms (i.e. sets of step-by-step instructions) to predict susceptibility to commonly-used antibiotics using routinely available patient data from a children’s hospital in Cambodia.
- We found that an algorithm called random forests enabled surprisingly accurate predictions, particularly for predicting whether the infection was likely to be treatable with ceftriaxone, the most commonly used empiric antibiotic at the study hospital.
- Using this approach it would be possible to correctly predict when a different antibiotic would be needed for empiric treatment over 80% of the time, while recommending a different antibiotic when ceftriaxone would suffice less than 20% of the time.

What Do These Findings Mean?
- Using readily available patient information, sophisticated algorithms can enable good predictions of whether antibiotics are likely to be effective several days before laboratory tests are available.
- Algorithms would need to be trained with local hospital data, but our study shows that even with relatively limited data from a small hospital, good predictions can be obtained.
- Used as part of a decision support system such algorithms could help choose appropriate antibiotics for empiric therapy; this would be expected to translate into better patient outcomes and may help to reduce resistance.
- Such as a decision support system would have very low costs and be easy to implement in low- and middle-income countries.

## Introduction

There is consistent evidence that early and appropriate treatment of sepsis can reduce mortality [1]. Since definitive identification of a bacterial pathogen and its antibiotic susceptibility typically takes three to four days using conventional culture methods, empiric antibiotic therapy (i.e. therapy that starts before the causative organism and its antibiotic susceptibility is known) is recommended. Choice of empirical antibiotic aims to balance two objectives: first, to cast a wide spectrum of coverage effective against the most likely causative organisms; second, to minimize the selection of resistance to reserve antibiotics in the wider population [2]. Balancing the consequences associated with these two concerns - immediate patient outcomes and long-term resistance patterns impacting on future patients - represents a major challenge.

Empiric antibiotic choice for invasive bacterial infections in hospitalized children in low-to-middle income countries (LMICs) constitutes a particularly stark example of this problem owing to the high attributable mortality [3], and the high prevalence of antimicrobial resistance, particularly in neonates [4].

Current World Health Organization (WHO) guidelines for suspected sepsis or serious bacterial infection in newborns recommend empirical usage of gentamicin and ampicillin as the first line therapy, and change to third-generation cephalosporins if there is lack of improvement in 24-72 hours. [5,6] However, a systematic review of community-acquired neonatal sepsis in developing countries in 2012 found that of the causative pathogens in older infants (1-12 months), only 63% and 64% showed *in vitro* susceptibility to ampicillin and gentamicin, and third-generation cephalosporins, respectively. [6] For neonates, susceptibilities were even lower, with only 57% and 56% of pathogens susceptible to ampicillin and gentamicin and third-generation cephalosporins, respectively.

The potential harms of widespread antimicrobial resistance in children were illustrated in a recent study performed between 2007 and 2016 in a Cambodian children’s hospital, which found those infected with third-generation cephalosporin-resistant bacteria were less likely than others to receive appropriate antimicrobial therapy (57% vs. 94%), and when appropriate therapy was administered, it was initiated later (2 days vs. 0 days after admission for those who survived; 0.5 days vs. 0 days for those who died) [7]. In multivariable logistic regression, third-generation cephalosporin resistance was independently associated with death (aOR 2.65, 95% CI 1.05-6.96); p = 0.042) and intensive care unit (ICU) admission (aOR 3.17, 95% CI 1.31-8.10).

While anticipated clinical efficacy is the primary deciding factor in empirical antibiotic choices, [8] there are other important considerations as well. These include side effect profile [9], cost, ease of administration and risks of promoting resistance emergence in hospital settings. [2]

The adoption of antimicrobial stewardship programmes in hospitals is widely advocated internationally. This is true both in LMICs and high income countries where they are increasingly deployed, but still have substantial potential for improvement [10,11]. Locally-adapted hospital antibiotic policies are important components of such programmes, and typically contain recommendations for empiric antibiotic use. In most cases these recommendations are derived from expert opinion and informal (non-quantitative) syntheses of available evidence [12]. In some cases simple decision support systems based on logistic regression models and score systems have been developed to help identify patients at high risk of being infected with multidrug-resistant pathogens. These approaches have primarily been developed in high- and upper middle-income countries [13-18]. The use of predictive modeling as part of clinical decision support systems for antimicrobial management remains rudimentary, with only one example identified in a recent systematic review [19]. It has, however, been demonstrated in a randomized trial (in Israel, Germany and Italy) that a computerized decision support systems making use of an underlying causal probabilistic network model can lead to more appropriate empiric antibiotic prescribing [20].

We hypothesized that applying modern machine learning approaches to readily collected patient data can surpass the performance of those based on logistic regression or simple decision trees, and derive patient-specific predictions for antibiotic susceptibility. Improved predictions directing empirical antibiotic therapy may contribute to better patient outcomes while avoiding the overuse of inappropriate antibiotics that select for resistance.

In this study, we propose a locally adapted decision support system for a Cambodian children’s hospital by applying an array of machine learning algorithms to patient-level data. We evaluated the ability of the algorithms to predict whether the causative organisms were susceptible to (i.e. treatable with): i) ampicillin and gentamicin; ii) ceftriaxone; iii) neither i) nor ii). We specifically focus on the value of using the predictive models to identify patients at high risk of being infected with organisms resistant to ceftriaxone, a third-generation cephalosporin and the most commonly prescribed empirical antibiotic in practice at our study site.

## Materials and methods

### Data

Retrospective data were collected from the Angkor Hospital for Children, a non-governmental hospital in Siem Reap, North-western Cambodia with approximately 100 beds, and its Satellite Clinic situated 30km away, with 20 inpatient beds. The hospital provides free surgical and general medical care to children less than 16 years of age and is equipped with an ICU. Admitted neonates and children come from both urban and rural settings, with about two thirds residing in Siem Reap province. Over 90% of patients admitted come from the community, the rest being transferred from another hospital or clinic. None of the children are born in the hospital as there is no obstetric service.

Blood cultures are routinely taken from febrile inpatients (axillary temperature > 37.5°C) in accordance with clinical algorithms. Processing of these cultures including *in vitro* antibiotic susceptibility testing has been described elsewhere [21]. Children with at least one positive blood culture for a bacterial pathogen (other than coagulase negative *Staphylococci* and other likely skin contaminants) isolated between February 2013 and January 2016 were included in this study. We considered routine clinical data collected by the hospital and data on living conditions including household size, presence of domestic animals, and factors relating to water and sanitation. The study was approved by the Angkor Hospital for Children Institutional Review Board (AHC-IRB, 290) and the Oxford Tropical Research Ethics Committee (OxTREC, 08-12).

### Data Analysis

We evaluated a suite of machine learning algorithms based on their ability to predict the invasive pathogens’ Gram stain and *in vitro* susceptibility to antibiotics using available information prior to receiving culture results. Specifically, the antibiotics considered were: i) ampicillin and gentamicin; ii) ceftriaxone; iii) either i) or ii). In the event that more than one organism was grown from the same blood culture, they were categorized as susceptible to the specified antibiotics only if all organisms were susceptible to at least one.

To predict the above four dependent variables we selected 35 independent variables (predictors) from patient records by coding quality and relevance. Dichotomous predictors where all but 10 or fewer patients had the same value were excluded. Missing data for binary predictors were treated as negative (e.g. NA for domestic animal was considered no domestic animal).

Weight for age standard deviations (z-score), a measure of malnutrition, was calculated using the LMS method [22] based on growth charts from the Centers for Disease Control. All dataset files are available at http://doi.org/10.5281/zenodo.1256967.

#### Training the algorithms

For comparison, a logistic regression with backwards step-wise AIC model selection was performed [23]. Additional machine learning algorithms that were explored include decision trees constructed via recursive partitioning [24], random forests [25], boosted decision trees using adaptive boosting [26], linear support vector machines (SVM) [27], polynomial SVMs, radial SVMs [28] and k-nearest neighbors [29]. All analysis was done in R [30] using the following packages; MASS [31] (stepwise logistic regression), rpart [32] (decision tress), ranger [33] (random forest), fastAdaboost [34] (boosted decision trees), kknn [35] (k-nearest neighbors), kernlab [36] (polynomial and radial SVM), and LiblineaR [37] (linear SVM). Parameters were fitted for highest Kappa based on a grid search [38]. Machine learning models were 5-fold cross-validated repeated 3 times.

An 80/20 split for training and testing data set was adopted. For categorical variables we ensured that each category was represented by at least one record in the training set. To assess algorithm performance over different training sets, each model was refitted to 100 random selections of training and testing data sets. Performance was ranked based on area under the receiver operating characteristic curve (AUC) from the test set.

For each method we report both the performance ranking for different outcomes, and then consider each method’s individual performance. We select the best method overall, then consider its probability calibration and the most important predictors. Variable importance in random forests was calculated using the method described in *Janitza et al*. [39]

#### Identifying the optimum cut-off

The Receiver Operating Characteristic (ROC) curves show what sensitivity (i.e. chance of correctly identifying a non-susceptible infection) each test would be expected to have for a given specificity (i.e chance of correctly identifying a susceptible infection). However, they do not tell us what optimal cut-off for specificity we should use in practice. One possible approach would be to choose this cut-off to maximize the overall test *accuracy* (i.e. the chance the test gives a true positive or true negative result). However, this would ignore the fact that the cost (in terms of economic loss and health impact) of failing to identify a non-susceptible infection will in general differ from the cost of mistakenly classifying a susceptible infection as non-susceptible. To identify the optimum cut-off, we should instead seek to choose the specificity that, for a given ROC curve, maximizes the utility. This can be accomplished by adopting a health economic framework, taking into account direct and indirect values of both economic costs and changes in health outcomes associated with different cut-offs. Costs include those of the prescribed antibiotics, excess length of stay and impact on mortality when no effective empiric antibiotic is prescribed, and, most challengingly, future impact on health outcomes due to selection for resistance that antibiotic usage may cause (i.e. whether the use is appropriate or not). Given that the cost of future resistance is difficult to quantify, an alternative approach is to consider willingness to pay (WTP) for avoiding unnecessary use of last-line (in this case carbapenem) antibiotics. With this economic framework, and using conventional recommendations for WTP per quality adjusted life year (QALY) gained [40], health impact and monetary costs can be combined on the same scale and represented as net monetary value (monetary loss + QALY loss x WTP). In this way we can assign different net monetary values to each of the four possible test outcomes (true positive, true negative, false positive, false negative). The optimal cut-off will be a value of the specificity that minimizes this net monetary loss. We provide illustrative examples of these calculations (see S1 Appendix for further details) and provide a user-friendly web application to enable optimal cut-offs to be determined under different assumptions, available at https://mathu.shinyapps.io/rf-ceft-cost/.

## Results

Fig 1 shows the selection of cases used for model training and testing. Of 245 cases, two cases were excluded; one due to missing target outcome data, and the other due to a biologically impossible value.

**Fig 1.**
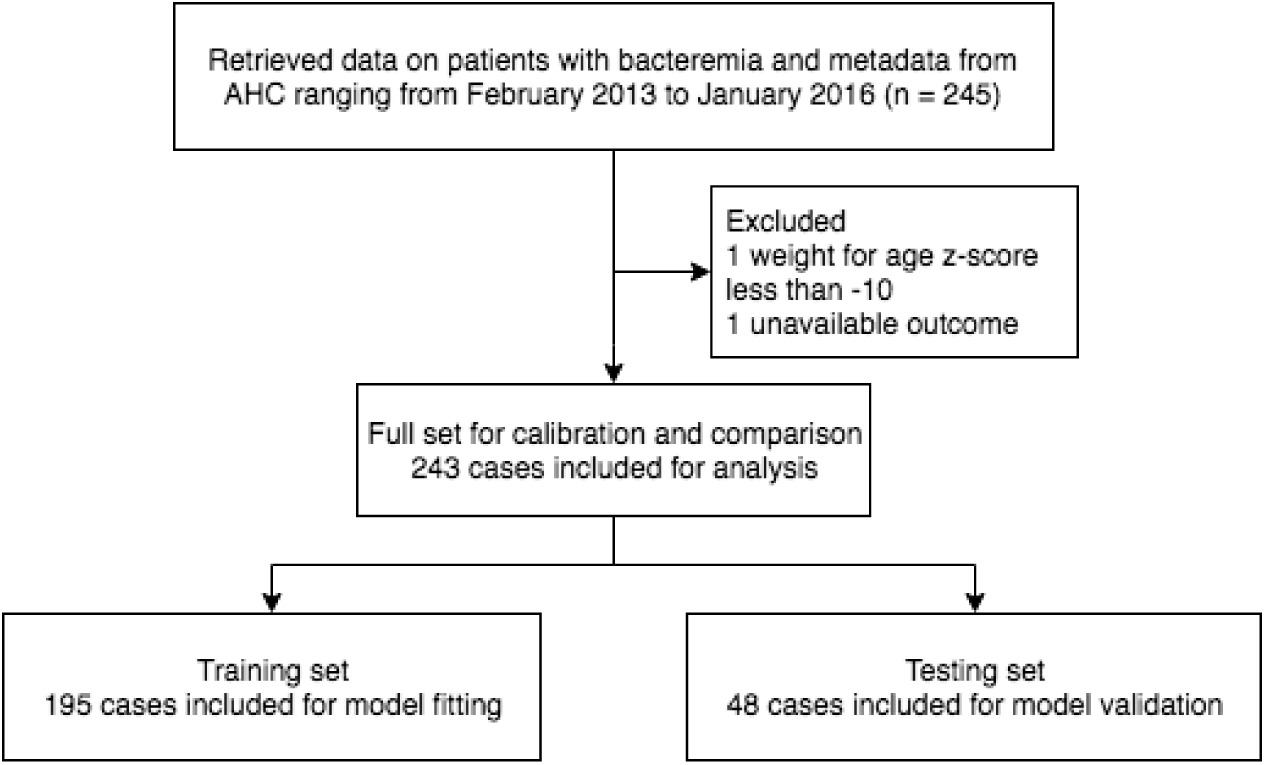
Selection of records.

Comparing relative performance of the approaches based on the AUC, the random forest method was most frequently ranked first, and was consistently ranked higher than decision trees, boosted decision trees, k-nearest neighbors, and the widely-used stepwise logistic regression. Of the four dependent variables, lack of susceptibility to ampicillin and gentamicin (Fig 2A), and to ampicillin, gentamicin and ceftriaxone (Fig 2C) were best predicted by the random forest approach.

**Fig 2.**
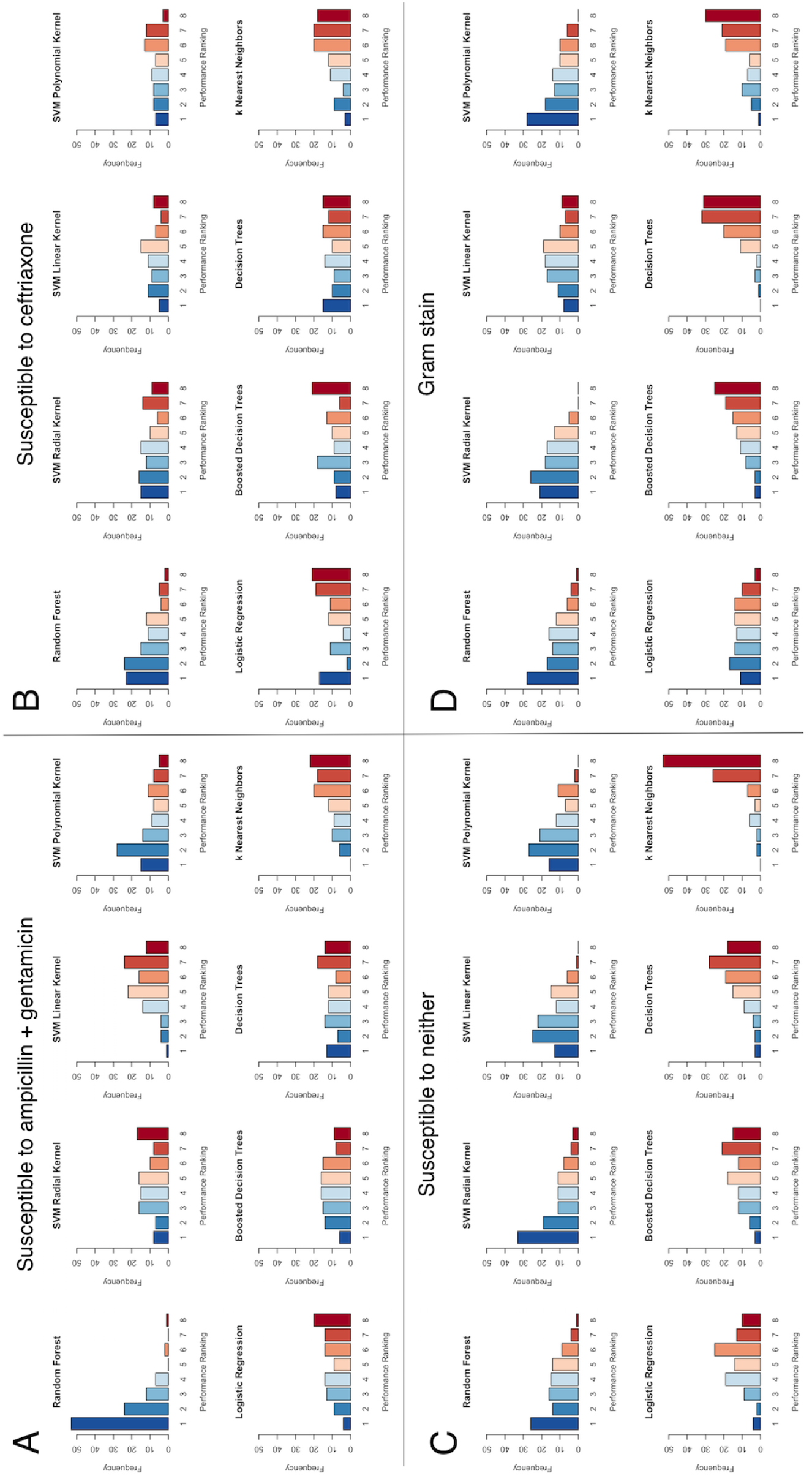
Comparison of performance rankings. Histograms of performance rankings obtained with 100 random splits of data into training (80%) and testing (20%) sets for the eight machine learning algorithms for predicting four outcomes (A) lack of susceptibility to ampicillin + gentamicin (B) lack of susceptibility to ceftriaxone (C) lack of susceptibility to both (D) Gram stain. Ranking of 1 (blue) is best, 8 (red) is worst. Rankings are based on the area under the ROC curves with the test data.

Overall, the SVM approaches performed well, but with some variation depending on which kernel the models were based on and which outcomes were being considered. For example, an SVM with radial kernel outperformed the random forest approach by a small margin in predicting lack of ceftriaxone susceptibility (Fig 2B), but performed quite poorly for predicting lack of susceptibility to ampicillin and gentamicin (Fig 2A). None of the three SVM variants performed well for all four of the outcomes. Even the algorithms that performed worst overall performed best for some of the randomly-selected training/test data splits. Thus, while the k-nearest neighbors method was most frequently ranked the lowest out of all the methods, for a small number of random selections of training data this algorithm was the best for predicting Gram stain and lack of susceptibility to all three antibiotics.

Ranking, although a good indicator of relative performance, does not necessarily indicate prediction ability itself. Fig 3 shows Receiver Operating Characteristic (ROC) curves for predicting lack of susceptibility to ceftriaxone for all methods. This figure highlights the disconnect between predictive performance on the training data set (blue dotted line) and that on the test set (black dashed lines), highlighting the importance of separating the training and testing data. It is possible for ROC curves for different methods to cross, indicating that optimal methods may vary depending on the cut-off used, and that the methods with the highest AUC may not always be the best for a given application. Importantly, the random forest test set ROC curve did not cross with other test set ROC curves.

**Fig 3.**
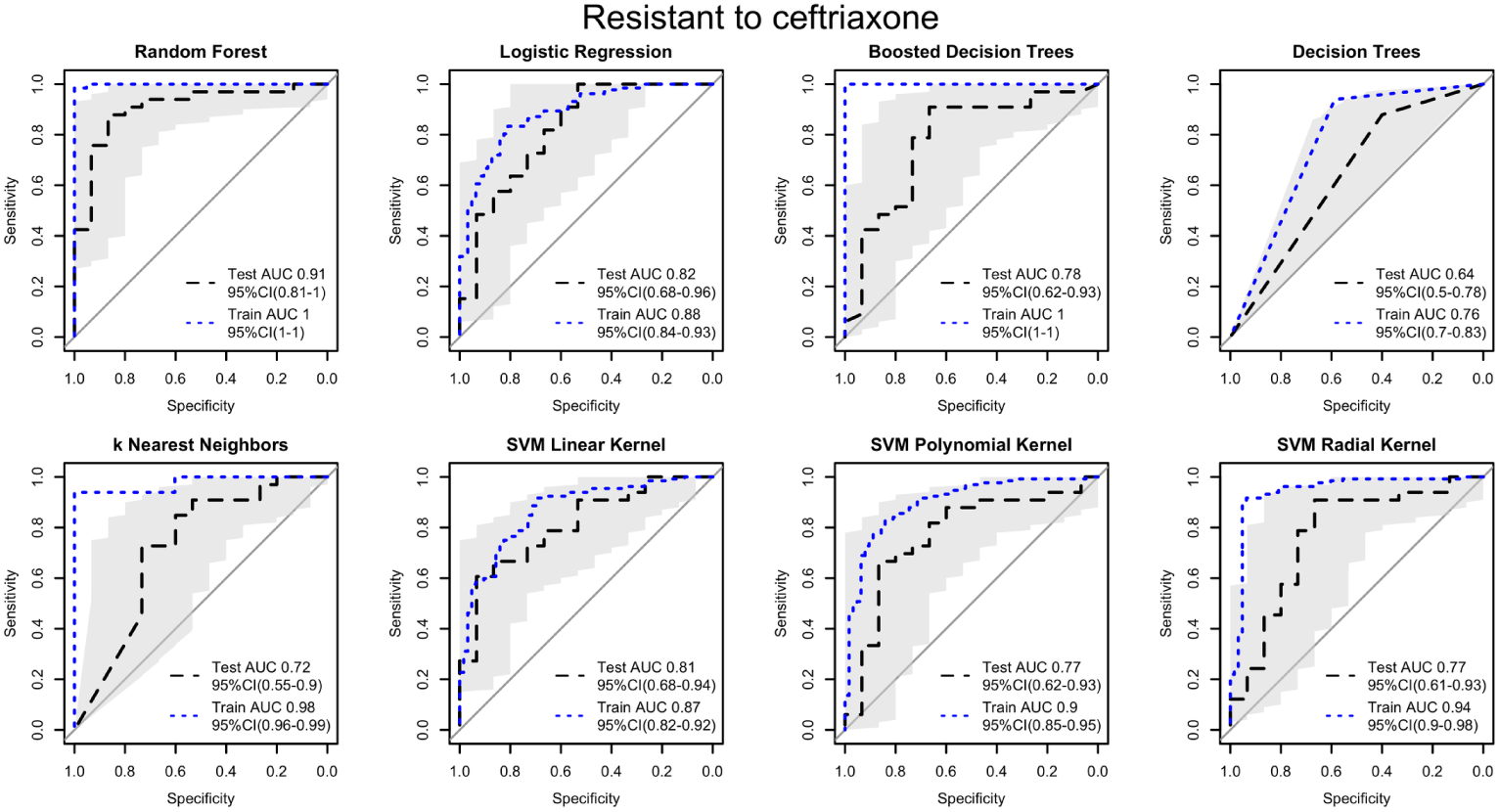
Receiver Operating Characteristic (ROC) curves for predicting lack of susceptibility to ceftriaxone. Training set (blue dotted lines), testing set i.e. actual performance (black dashed lines with 95% intervals shown by shading). The solid diagonal line is the line of no-discrimination, the expected performance of a random guess.

To be effective in supporting decisions, it is useful to not only rank well (predict correctly), but also to be well-calibrated (i.e. the estimated probabilities that pathogens lack susceptibility to an antibiotic should be similar to observed frequencies). A calibration plot for the ceftriaxone outcome with random forests algorithm is shown in Fig 4 after Platt scaling of predicted probabilities [41]. This indicates that the model is well-calibrated.

**Fig 4.**
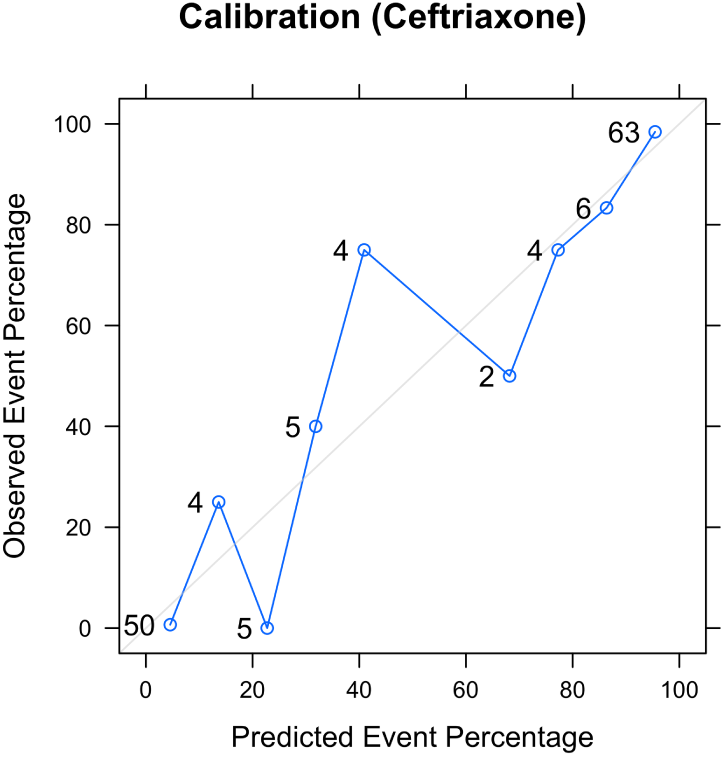
Calibration for random forest predicting lack of susceptibility to ceftriaxone. This compares predicted probabilities (grouped into 10 equally-sized bins) to observed event frequencies in real data using the entire data set. Points close to the gray diagonal line indicate that the predicted probability is close to the observed frequency. Numbers above points indicate the number of records contributing to each point.

Fig 5 illustrates the influence of each independent variable on the random forest model in predicting antibiotic susceptibilities. The most important predictor was days from hospital admission to blood sample, which would decrease the impurity of tree splits 100% of the time. That is to say, when the model was retrained with values from days from hospital admission permuted, it had a negative effect on the model’s ability to predict on all permutations. The assumption here is that if a predictor is important to our model, our model should be sensitive to changes from it. Changes were permuted from existing data to add realism to its range of possible values. In terms of importance, Days from hospital admission to blood sample was closely followed by the patient’s age, the classification of the infection as hospital- or community-acquired, and the patient’s weight (adjusted for age), which perturbations had an effect 75% of the time. For ceftriaxone, changes to variables related to previous hospital exposure and living conditions all only had an effect on the model less than 25% of the time.

**Fig 5.**
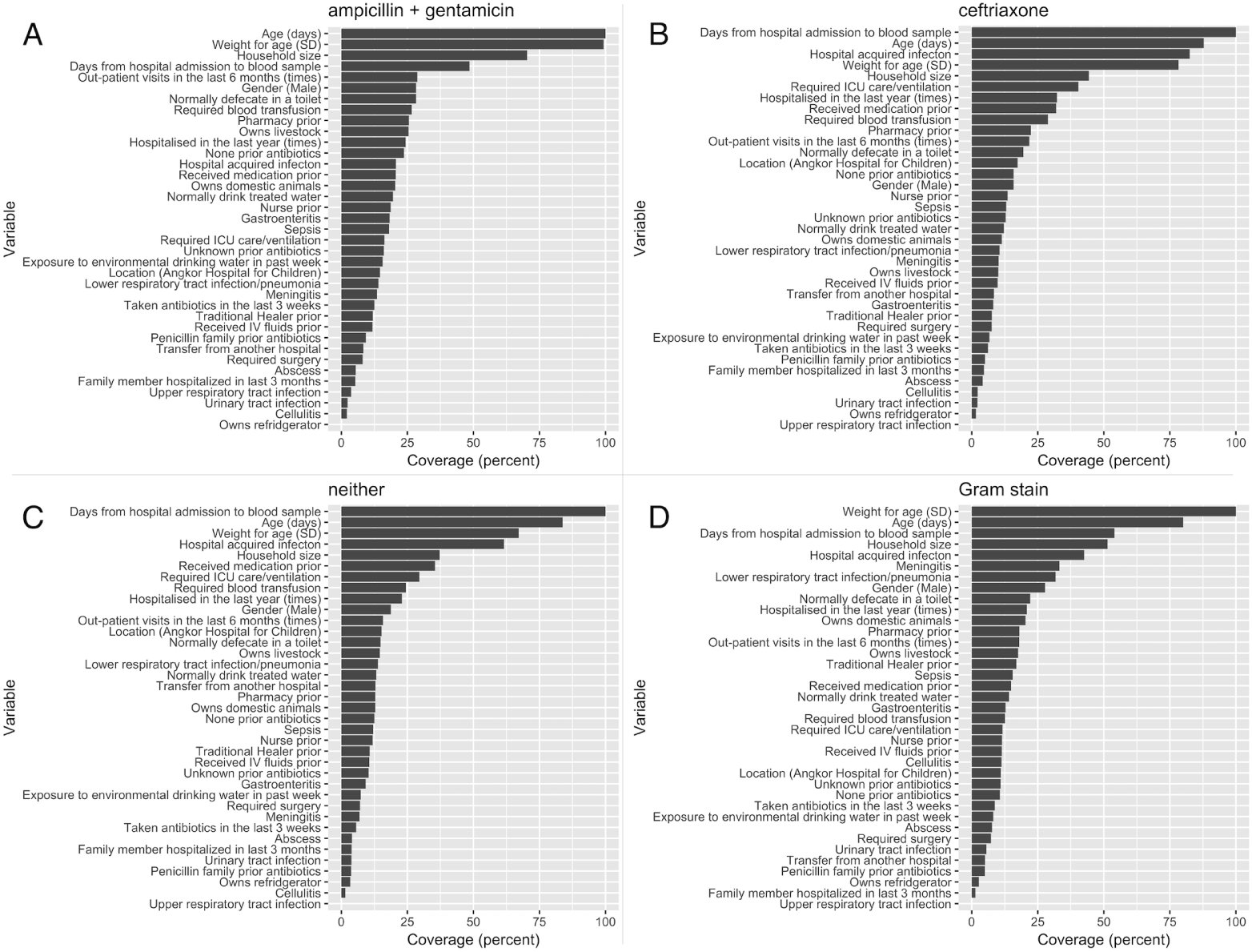
Variable importance in random forest models. Results show relative importance of variables for predicting lack of susceptibilty to ampiciliin + gentamicin (A) lack of susceptibilty to ceftriaxone (B) lack of susceptibility to both (C) Gram stain (D).

The most important predictors in the random forest model for the other three outcomes were broadly similar. Interestingly, the classification of infection as hospital- or community-acquired had less importance for predicting lack of susceptibility to ampicillin and gentamicin compared to ceftriaxone, but household size was found to much more important.

For comparison, in the widely-used stepwise-selected logistic regression model, important predictors in multivariable models for treatability by ceftriaxone were age, household size, meningitis, and hospital-versus community-acquired infection (Table 1).

With the exception of meningitis, all these predictors ranked high in coverage (top five) on importance in the random forests model.

Fig 6 illustrates how, used as part of a decision support system, choice of the test threshold to inform antibiotic prescribing decisions would impact on the number of patients treated empirically with appropriate antibiotics. Taking a test threshold of 0.29 for the predicted probability that ceftriaxone would not be an effective treatment (so above this value, patients would be recommended to receive a second-line antibiotic, typically a carbapenem, instead of ceftriaxone), 14 out of 15 (93%) patients in the test data set who have ceftriaxone-resistant infections would be correctly identified (true positives). This threshold choice would also lead to eight of the 33 (24%) patients with ceftriaxone-susceptible infections unnecessarily receiving the second-line antibiotic (over-treatment). Adjusting the threshold corresponds to moving the red line in Fig 6A-B up and down, changing the numbers of patients over- and under-treated. The choice of this threshold has an impact on patient outcomes and costs; their combined impact can be represented as the net utility loss (expressed as a net monetary value) due to infection (Fig 6D). A rational approach would be to choose the threshold to minimize this utility loss. However, quantifying utility loss due to future selection for resistance when using antibiotics is challenging [42], so an alternative approach is to choose a prediction threshold based on clinical judgment, and work backwards to determine how this implicitly values the utility loss due to over-treatment. In this example, we find a threshold of 0.29 implies that we would be willing to pay $US 200 to avoid one unnecessary course of a carbapenem. Details of the calculations can be found in the supplementary text.

**Fig 6.**
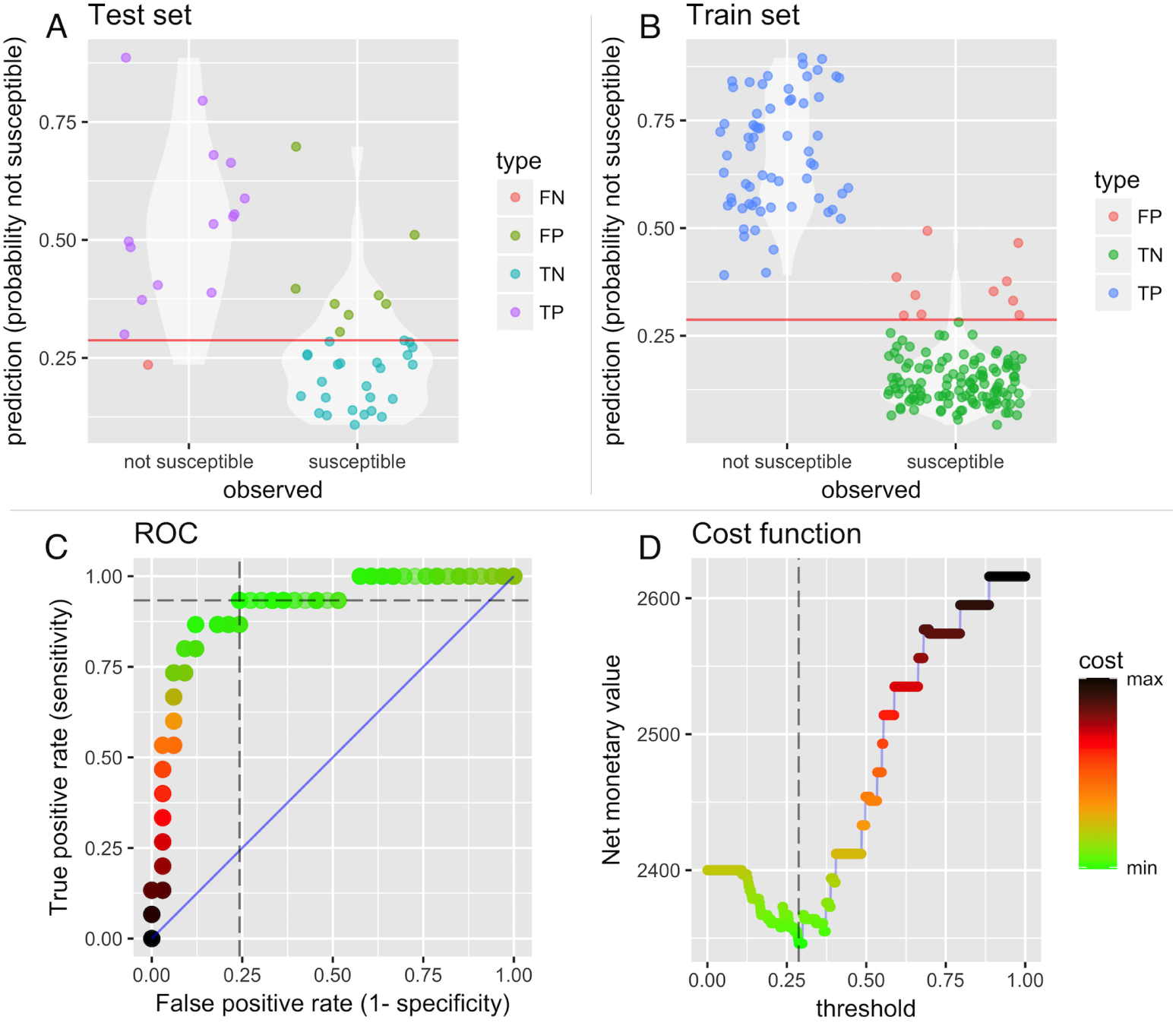
Effects of test cut-off (threshold) choice for decision outcomes and utility. Impact of test threshold (horizontal red line in panels A and B) on classification outcomes for lack of susceptibility to ceftriaxone, showing observations which, for the illustrated cut-off, are false negatives (FN), false positives (FP), true negatives (TN) and true positives (TP) in test (A) and training (B) data. The ROC curve (C) is shaded according to utility loss at different cut-offs, where horizontal dashed lines correspond to the threshold selected by minimizing the cost function (D), i.e. maximising utility. Higher utilities i.e. lower costs (expressed as a net monetary value) are shaded in green.

## Discussion

Our results show that modern machine learning algorithms can reliably and substantially outperform widely-used logistic regression models and provide accurate, actionable, and well-calibrated predictions about whether commonly used empirical antibiotics are likely to be effective. We found that the random forest approach performed particularly well, especially for predicting whether pathogens were likely to lack susceptibility to ceftriaxone, the most widely used empiric antibiotic for our study patients. To our knowledge this is the first time such machine learning algorithms have been applied to this problem.

The most important variables for predicting antibiotic susceptibility were found to be time from admission to blood culture, patient age, age-adjusted weight score, and hospital versus community-acquired infection. These are objective and routinely collected variables available in most clinical settings. All other variables included in the models are also easily collected at minimal cost through short questionnaires. The computations underlying the predictions can readily be performed in a few seconds on a low-cost computer, or remotely via any device connected to the Internet. This makes the approach highly suitable for other LMIC settings, which typically face the highest disease burden and the most urgent problems with antimicrobial resistance [43].

### Wider implications

Used as part of a decision support system, the best machine learning approaches should, in theory, make it possible to substantially increase the proportion of patients who receive effective empiric antibiotics, while minimizing the risks of increased resistance selection that would be associated with a blanket change in the default choice of empiric antibiotics for all patients. Clearly, further work is needed to evaluate such deployment in practice.

Rapid microbiological diagnostic tests offer an alternative pathway for improving the precision of early antibiotic prescribing. Affordable and accurate tests are not currently available, but this situation may change in the coming years. While machine learning approaches as proposed here could be considered a stopgap, we think it is more likely that the two approaches will be complementary. Results from future rapid diagnostic tests could be used as inputs into machine learning algorithms along with other patient variables, and would be expected to lead to more reliable predictions than those from the rapid tests alone.

**Table 1.** Distribution of variables for logistic regression for susceptibility to ceftriaxone.

| Characteristics | Treatable<br>n = 127<br>(No/Yes) | Resistant<br>n = 68<br>(No/Yes) | OR | 95% CI | P-value |
| --- | --- | --- | --- | --- | --- |
| Age (days) | 703; 1063* | 1616; 1613* | 1.205 | 1.032-1.41 | 0.0183 |
| Complication during admission |  |  |  |  |  |
| Required ICU care/ventilation | 32/31 | 23/109 | 0.767 | 0.203-2.89 | 0.6952 |
| Required surgery | 7/56 | 10/122 | 0.740 | 0.138-3.98 | 0.7260 |
| Required blood transfusion | 28/35 | 27/105 | 0.579 | 0.190-1.77 | 0.3372 |
| Transfer from another hospital | 18/45 | 18/114 | 0.295 | 0.080-1.09 | 0.0670 |
| Admission differential diagnosis |  |  |  |  |  |
| Sepsis | 37/26 | 82/50 | 0.537 | 0.174-1.66 | 0.2796 |
| Meningitis | 4/59 | 23/109 | 11.884 | 1.577-89.57 | 0.0163 |
| Lower respiratory tract infection/pneumonia | 17/46 | 33/99 | 0.556 | 0.180-1.72 | 0.3081 |
| Upper respiratory tract infection | 3/60 | 6/126 | 0.601 | 0.068-5.33 | 0.6472 |
| Gastroenteritis | 9/54 | 18/114 | 1.777 | 0.399-7.91 | 0.4508 |
| Cellulitis | 4/59 | 11/121 | 0.959 | 0.146-6.30 | 0.9650 |
| Abscess | 2/61 | 10/122 | 1.779 | 0.243-13.04 | 0.5708 |
| Urinary tract infection | 2/61 | 12/120 | 0.555 | 0.075-4.10 | 0.5643 |
| Weight for age (SD) | -2.2; 1.7* | -2.1; 1.7* | 1.322 | 0.990-1.76 | 0.0588 |
| Hospitalised in the last year (times) | 0; 0-3‡ | 0; 0-3‡ | 0.425 | 0.168-1.08 | 0.0719 |
| Out-patient visits in the last 6 months (times) | 0; 0-3‡ | 0; 0-3‡ | 0.873 | 0.410-1.86 | 0.7257 |
| Treatment prior to current admission |  |  |  |  |  |
| Pharmacy | 8/55 | 43/89 | 1.841 | 0.488-6.94 | 0.3674 |
| Nurse | 22/41 | 64/68 | 1.301 | 0.422-4.01 | 0.6472 |
| Traditional Healer (Khru Khmer) | 8/55 | 15/117 | 1.195 | 0.224-6.39 | 0.8347 |
| Received IV fluids | 11/52 | 31/101 | 0.771 | 0.182-3.26 | 0.7234 |
| Received medication | 34/29 | 105/27 | 0.699 | 0.161-3.03 | 0.6322 |
| Household size | 6; 3-10‡ | 6; 3-10‡ | 1.301 | 1.003-1.69 | 0.0472 |
| Owens domestic animals | 49/14 | 92/40 | 0.551 | 0.152-2.00 | 0.3649 |
| Owens livestock | 44/19 | 89/43 | 0.329 | 0.096-1.12 | 0.0762 |
| Normally defecate in a toilet | 33/30 | 62/70 | 0.504 | 0.179-1.42 | 0.1959 |
| Owens refridgerator | 4/59 | 5/127 | 0.852 | 0.090-8.04 | 0.8890 |
| Taken antibiotics in the last 3 weeks | 4/59 | 22/110 | 8.152 | 0.738-90.07 | 0.0869 |
| Family member hospitalized in last 3 months | 5/58 | 9/123 | 0.429 | 0.079-2.33 | 0.3266 |
| Exposure to environmental drinking water in past week | 7/56 | 24/108 | 0.242 | 0.050-1.18 | 0.0793 |
| Normally drink treated water | 26/37 | 61/71 | 1.071 | 0.391-2.93 | 0.8944 |
| Hospital acquired infecton | 34/29 | 11/121 | 0.067 | 0.013-0.34 | 0.0011 |
| Days from hospital admission to blood sample | 0; 0-104‡ | 0; 0-104‡ | 0.991 | 0.914-1.07 | 0.8308 |
| Gender (Male) | 33/30 | 79/53 | 0.684 | 0.254-1.84 | 0.4519 |
| Location (Angkor Hospital for Children) | 54/9 | 94/38 | 0.352 | 0.091-1.35 | 0.1282 |
| Taken antibiotics prior to admission |  |  |  |  |  |
| None | 39/24 | 62/70 | 51108763.849 | 0.000-Inf | 0.9882 |
| Penicillin Family | 4/59 | 17/115 | 68311352.311 | 0.000-Inf | 0.9880 |
| Unknown | 17/46 | 53/79 | 87658692.420 | 0.000-Inf | 0.9878 |

\*Mean; SD for normal distributions, ‡Mode; Range for exponential distributions,
SD, standard deviation; CI, confidence interval; OR, odds ratio; Inf, infinity
Odds ratio from multivariate logistic regression analysis prior to step-wise backward elimination

### Utility

A common dilemma in designing diagnostic systems is to identify the optimum cut-off point for sensitivity and specificity on the ROC curve. Increasing the sensitivity threshold for detecting lack of susceptibility to an antibiotic (which corresponds to lowering the horizontal red line in Fig 6) will capture more true cases of antimicrobial resistance where a reserve antibiotic (e.g. carbapenem) is appropriate and potentially improve patient outcomes (i.e. there will be more true positives). However, this will inadvertently lead to more prescriptions of reserve antibiotics when they are not needed (false positives), creating increased selection for resistance to what may be an antibiotic of last resort. Conversely, setting the threshold at higher specificity (raising the red line in Fig 6) has the benefit of reducing false positives, but the model will miss more patients with resistant bacterial infections (i.e. false negatives will increase), leading to delayed prescription of appropriate antibiotics. A natural approach would be to choose the cut-off to maximise utility (which includes health outcomes and opportunity costs associated with economic costs). While quantifying the direct healthcare cost components is relatively straightforward, the costs of resistance are far more challenging to calculate. *Shrestha et al* estimated the costs of resistance per antibiotic consumed, assigning a cost of $US 0.8 and $US 1.5 per standard unit of carbapenem in Thai and US settings, respectively [42]. However, these estimates did not take into account the potentially grave future consequences of losing an antibiotic of last resort. Better quantification of how we should value not using antibiotics is an important area of future research.

### Strengths and limitations

We systematically evaluated a number of machine learning algorithms to determine the algorithms with the best predictive performance for the problem. Most currently available clinical scoring systems rely on logistic regression models, probably for historical reasons. No method is universally better than another method [44,45], however different algorithms have strengths and weaknesses, and our results suggest that by focusing on a single learning algorithm much of the previous literature may have missed an important opportunity.

A second important strength of our work is that algorithm training and evaluation were performed on different data sets. As is clear from Fig 3, if this is not done inflated performance estimates are likely. Though there are some notable exceptions [13, 15, 18], this separation has not always been performed in the previous literature, and this would be expected to lead to predictive power that is substantially lower than reported.

Thirdly, our analysis maximizes accuracy towards local usage. If we had used a large dataset aggregated from multiple settings in the hope of increasing generalisability, algorithm performance for all of the contributing centres would have been likely to suffer. Generally, scoring systems developed in one setting have been found to have substantially worse performance in different settings [14,46]. Our approach focuses on generalizing towards new samples obtained within the same setting with a moderately-sized dataset by comparing a large number of models [47]. Importantly, this suggests that the potential benefits of machine learning, which are often assumed to depend on large high-quality data-sets more commonly available in high-income settings [48], may be considerable even in resource- and relatively data-poor settings.

There are several limitations to our study. As with most clinical predictive systems, generalizability is a concern. A model developed using data from one hospital, may have poor predictive value when applied in another setting [14, 46]. We anticipate that wider deployment of such approaches would require models to be tailored to local data. The model may also become less relevant as time passes. Identifying the most appropriate temporal and spatial selection windows for training data is an important area for future research.

### Understanding the algorithms

One potential obstacle to the wider adoption of machine learning algorithms is that, to many, they are a black box. An intuitive way to understand them is to consider a geometric interpretation, visualizing a smaller problem first. Suppose we have a dataset with two predictors, height and weight. We can imagine each data point inhabiting a point in a 2-dimensional plane, *feature space* (i.e. a graph with weight on the x-axis and height on the y-axis). Each point would have a label of the class we are trying to predict (i.e. diseased/healthy). A classification problem can be superficially phrased as a search to find a line (or lines) which best separates the points with different labels on its feature space. For example, a line which splits between disease and healthy people on the height-weight plane. These lines do not necessarily need to be straight. For two independent variables this can be visualized as a graph. For *n* predictors this would require *n*-dimensions. For *n* > 3 this is harder to visualize, but the geometric interpretation still holds.

A geometric visualization allows us to appreciate the varying performances of each method by considering how each method arrives at the conclusion as to which line (or combination of lines) is best. A decision tree can be considered a combination of decisions, each represented by a line in our feature plane (i.e. *is weight > 50 kg?* can be considered a line at 50 on the weight axis). A combination of simple lines allows for more complex decision boundaries. But because of their ability to create complex boundaries, they tend to overfit. Random forests are designed to correct for the habit of decision trees to overfit by building a consensus of a multitude of decision trees, and averaging these trees by giving the majority vote after polling all component decision trees based on classification. In contrast, *k*-nearest neighbor, our least effective model, simply considers classification based solely on the assumption that points closer in feature space should be given the same classification. SVM with a radial kernel, in contrast, looks for the circle that best separates points with different outcome labels. Our results (Fig 2) clearly show that chance plays an important role in determining which lines best separate the points (and therefore which algorithm wins), but some algorithms (random forests and SVM) tend to have a much better chance of doing well.

## Conclusion

Decision support systems, informed by analysis of readily available data, when calibrated to local data, have the potential to lead to evidence-based hospital antibiotic policies which could improve the chances patients receive the most appropriate empiric antibiotics. This would be expected to lead to better patient outcomes and could help minimize the risk of increasing antibiotic resistance. While guidelines for developing a hospital antibiotic policy advocate conducting literature reviews and basing recommendations on local cumulative surveillance antibiograms [12], we have shown that machine learning algorithms informed by relatively small amounts of patient-level data can be used for deriving targeted and well-calibrated patient specific predictions for what empirical antibiotic therapy is likely to be appropropriate. Such a prediction system can be developed cheaply, using easily-collected data, and is well-suited to LMIC settings.

## Acknowledgments

We’d like to thank the staff and patients at Angkor Hospital for Children for their invaluable contribution and support that made this work possible.

## Supporting information

**S1 Fig. ROC comparison for predicting for treatability by neither (amp+gent nor ceft)** blue dotted lines: training set, black dashed lines: testing test (actual performance)

**S2 Fig. ROC comparison for predicting for treatability by ampicillin + gentamicin** blue dotted lines: training set, black dashed lines: testing test (actual performance)

**S3 Fig. ROC comparison for predicting for gram-positive stain** blue dotted lines: training set, black dashed lines: testing test (actual performance)

**S4 Appendix. Economic model to identify the optimum cut off**

**S5 Table. Distribution of variables for logistic regression for treatability by ampicillin + gentamicin**

**S6 Table. Distribution of variables for logistic regression for treatability by neither** neither amp+gent nor ceft

**S7 Table. Distribution of variables for logistic regression for predicting gram-positive stain**

## References

1. Liu VX, Fielding-Singh V, Greene JD, Baker JM, Iwashyna TJ, Bhattacharya J, et al. The Timing of Early Antibiotics and Hospital Mortality in Sepsis. American Journal of Respiratory and Critical Care Medicine. 2017;196(7):856–863. doi:10.1164/rccm.201609-1848OC.

2. De Man P, Verhoeven B, Verbrugh H, Vos M, Van den Anker J. An antibiotic policy to prevent emergence of resistant bacilli. The Lancet. 2000;355(9208):973–978.

3. Liu L, Oza S, Hogan D, Chu Y, Perin J, Zhu J, et al. Global, regional, and national causes of under-5 mortality in 2000-15: an updated systematic analysis with implications for the Sustainable Development Goals. The Lancet. 2016;388(10063):3027–3035.

4. Lubell Y, Ashley EA, Turner C, Turner P, White NJ. Susceptibility of community-acquired pathogens to antibiotics in Africa and Asia in neonates-an alarmingly short review. Tropical Medicine & International Health. 2011;16(2):145–151.

5. Organization WH. Pocket book of hospital care for children: guidelines for the management of common childhood illnesses. World Health Organization; 2013.

6. Downie L, Armiento R, Subhi R, Kelly J, Clifford V, Duke T. Community-acquired neonatal and infant sepsis in developing countries: efficacy of WHO’s currently recommended antibiotics—systematic review and meta-analysis. Archives of disease in childhood. 2012; p. archdischild-2012.

7. Fox-Lewis A, Takata J, Miliya T, Lubell Y, Soeng S, Sar P, et al. Antimicrobial resistance in invasive bacterial infections in hospitalized children, Cambodia, 2007-2016. Emerging infectious diseases. 2018;24(5):841.

8. Mtitimila EI, Cooke RW. Antibiotic regimens for suspected early neonatal sepsis. The Cochrane library. 2004;.

9. Fuchs A, Zimmermann L, Graz MB, Cherpillod J, Tolsa JF, Buclin T, et al. Gentamicin exposure and sensorineural hearing loss in preterm infants. PloS one. 2016;11(7):e0158806.

10. Johannsson B, Beekmann SE, Srinivasan A, Hersh AL, Laxminarayan R, Polgreen PM, et al. Improving Antimicrobial Stewardship The Evolution of Programmatic Strategies and Barriers. Infection Control & Hospital Epidemiology. 2011;32(4):367–374.

11. Hersh AL, Beekmann SE, Polgreen PM, Zaoutis TE, Newland JG. Antimicrobial stewardship programs in pediatrics. Infection Control & Hospital Epidemiology. 2009;30(12):1211–1217.

12. Organization WH. Step-by-step approach for development and implementation of hospital antibiotic policy and standard treatment guidelines; 2011.

13. Tumbarello M, Trecarichi EM, Bassetti M, De Rosa FG, Spanu T, Di Meco E, et al. Identifying patients harboring extended-spectrum-β-lactamase-producing Enterobacteriaceae on hospital admission: derivation and validation of a scoring system. Antimicrobial agents and chemotherapy. 2011;55(7):3485–3490.

14. Pan A, Lee A, Cooper B, Chalfine A, Daikos G, Garilli S, et al. Risk factors for previously unknown meticillin-resistant Staphylococcus aureus carriage on admission to 13 surgical wards in Europe. Journal of Hospital Infection. 2013;83(2):107–113.

15. Lee AS, Pan A, Harbarth S, Patroni A, Chalfine A, Daikos GL, et al. Variable performance of models for predicting methicillin-resistant Staphylococcus aureus carriage in European surgical wards. BMC infectious diseases. 2015;15(1):105.

16. Kengkla K, Charoensuk N, Chaichana M, Puangjan S, Rattanapornsompong T, Choorassamee J, et al. Clinical risk scoring system for predicting extended-spectrum β-lactamase-producing Escherichia coli infection in hospitalized patients. Journal of Hospital Infection. 2016;93(1):49–56. doi:10.1016/j.jhin.2016.01.007.

17. Johnson SW, Anderson DJ, May DB, Drew RH. Utility of a clinical risk factor scoring model in predicting infection with extended-spectrum β-lactamase-producing enterobacteriaceae on hospital admission. Infection Control & Hospital Epidemiology. 2013;34(4):385–392.

18. Tumbarello M, Trecarichi EM, Tumietto F, Del Bono V, De Rosa FG, Bassetti M, et al. Predictive models for identification of hospitalized patients harboring KPC-producing Klebsiella pneumoniae. Antimicrobial agents and chemotherapy. 2014; p. AAC-02373.

19. Rawson T, Moore L, Hernandez B, Charani E, Castro-Sanchez E, Herrero P, et al. A systematic review of clinical decision support systems for antimicrobial management: are we failing to investigate these interventions appropriately? Clinical Microbiology and Infection. 2017;23(8):524–532.

20. Paul M, Andreassen S, Tacconelli E, Nielsen AD, Almanasreh N, Frank U, et al. Improving empirical antibiotic treatment using TREAT, a computerized decision support system: cluster randomized trial. Journal of Antimicrobial Chemotherapy. 2006;58(6):1238–1245.

21. Stoesser N, Moore CE, Pocock JM, An KP, Emary K, Carter M, et al. Pediatric bloodstream infections in Cambodia, 2007 to 2011. The Pediatric infectious disease journal. 2013;32(7):e272–e276.

22. Cole TJ. The LMS method for constructing normalized growth standards. European journal of clinical nutrition. 1990;44(1):45–60.

23. Venables WN, Ripley BD. Modern Applied Statistics with S. 4th ed. New York: Springer; 2002. Available from: http://www.stats.ox.ac.uk/pub/MASS4.

24. Breiman L, Friedman J, Stone CJ, Olshen RA. Classification and Regression Trees. The Wadsworth and Brooks-Cole statistics-probability series. Taylor & Francis; 1984. Available from: https://books.google.co.th/books?id=JwQx-WOmSyQC.

25. Breiman L. Random Forests. Machine Learning. 2001;45(1):5–32. doi:10.1023/A:1010933404324.

26. Freund Y, Schapire RE, et al. Experiments with a new boosting algorithm. In: Icml. vol. 96. Bari, Italy; 1996. p. 148–156.

27. Hearst MA, Dumais ST, Osuna E, Platt J, Scholkopf B. Support vector machines. IEEE Intelligent Systems and their applications. 1998;13(4):18–28.

28. Scholkopf B, Smola AJ. Learning with kernels: support vector machines, regularization, optimization, and beyond. MIT press; 2001.

29. Mitchell TM, et al. Machine learning. 1997. Burr Ridge, IL: McGraw Hill. 1997;45(37):870–877.

30. R Core Team. R: A Language and Environment for Statistical Computing; 2016. Available from: https://www.R-project.org/.

31. Venables WN, Ripley BD. Modern Applied Statistics with S. 4th ed. New York: Springer; 2002. Available from: http://www.stats.ox.ac.uk/pub/MASS4.

32. Therneau T, Atkinson B, Ripley B. rpart: Recursive Partitioning and Regression Trees; 2017. Available from: https://CRAN.R-project.org/package=rpart.

33. Wright MN, Ziegler A. ranger: A Fast Implementation of Random Forests for High Dimensional Data in C++ and R. Journal of Statistical Software. 2017;77(1):1–17. doi:10.18637/jss.v077.i01.

34. Chatterjee S. fastAdaboost: a Fast Implementation of Adaboost; 2016. Available from: https://CRAN.R-project.org/package=fastAdaboost.

35. Schliep K, Hechenbichler K. kknn: Weighted k-Nearest Neighbors; 2016. Available from: https://CRAN.R-project.org/package=kknn.

36. Karatzoglou A, Smola A, Hornik K, Zeileis A. kernlab – An S4 Package for Kernel Methods in R. Journal of Statistical Software. 2004;11(9):1–20.

37. Helleputte T. LiblineaR: Linear Predictive Models Based on the LIBLINEAR C/C++ Library; 2017.

38. McHugh ML. Interrater reliability: the kappa statistic. Biochemia Medica. 2012;22(3):276–282.

39. Janitza S, Celik E, Boulesteix AL. A computationally fast variable importance test for random forests for high-dimensional data. Advances in Data Analysis and Classification. 2016; p. 1–31.

40. Organization WH, et al. Macroeconomics and health: investing in health for economic development: report of the Commission on Macroeconomics and Health. In: Macroeconomics and health: investing in health for economic development: report of the commission on macroeconomics and health; 2001.

41. Niculescu-Mizil A, Caruana R. Predicting Good Probabilities with Supervised Learning. In: Proceedings of the 22Nd International Conference on Machine Learning. ICML ’05. New York, NY, USA: ACM; 2005. p. 625–632. Available from: http://doi.acm.org/10.1145/1102351.1102430.

42. Shrestha P, Cooper BS, Coast J, Oppong R, Do NTT, Podha T, et al. Enumerating the Economic Cost of Antimicrobial Resistance Per Antibiotic Consumed to Inform the Evaluation of Interventions Affecting their Use. bioRxiv. 2017;doi:10.1101/206656.

43. Okeke IN, Laxminarayan R, Bhutta ZA, Duse AG, Jenkins P, O’Brien TF, et al. Antimicrobial resistance in developing countries. Part I: recent trends and current status. The Lancet infectious diseases. 2005;5(8):481–493.

44. Wolpert DH, Macready WG. No free lunch theorems for optimization. IEEE transactions on evolutionary computation. 1997;1(1):67–82.

45. Caruana R, Niculescu-Mizil A. An empirical comparison of supervised learning algorithms. In: Proceedings of the 23rd international conference on Machine learning. ACM; 2006. p. 161–168.

46. Slekovec C, Bertrand X, Leroy J, Faller JP, Talon D, Hocquet D. Identifying patients harboring extended-spectrum-β-lactamase-producing Enterobacteriaceae on hospital admission is not that simple. Antimicrobial agents and chemotherapy. 2012;56(4):2218–2219.

47. Shaikhina T, Lowe D, Daga S, Briggs D, Higgins R, Khovanova N. Machine learning for predictive modelling based on small data in biomedical engineering. IFAC-PapersOnLine. 2015;48(20):469–474.

48. Salathé M. Digital epidemiology: what is it, and where is it going? Life sciences, society and policy. 2018;14(1):1.

